# Developing and adult reef fish show rapid, reversible light-induced plasticity in their visual system

**DOI:** 10.1101/2022.06.02.494615

**Authors:** Lily G. Fogg, Fabio Cortesi, Camille Gache, David Lecchini, N. Justin Marshall, Fanny de Busserolles

**Affiliations:** Queensland Brain Institute, The University of Queensland, Brisbane, Queensland, 4072, Australia; PSL Research University, EPHE-UPVD-CNRS, UAR3278 CRIOBE, 98729 Papetoai, Moorea, French Polynesia; Laboratoire d’Excellence “CORAIL”, Paris, France

**Keywords:** Retina, Vision, Teleost fish, Phenotypic plasticity, Light environment, Ontogeny

## Abstract

The visual capabilities of fish are optimised for their ecology and light environment over evolutionary time. Similarly, fish vision can adapt to regular changes in light conditions within their lifetime, *e*.*g*., ontogenetic or seasonal variation. However, we do not fully understand how vision responds to irregular short-term changes in the light environment, *e*.*g*., algal blooms and light pollution. In this study, we investigated the effect of short-term exposure to unnatural light conditions on opsin gene expression and retinal cell densities in larval and adult diurnal reef fish (convict surgeonfish; *Acanthurus triostegus*). Results revealed phenotypic plasticity in the retina across ontogeny, particularly in the larvae. The most substantial differences at both molecular and cellular levels were found under constant dim light, while constant bright light or simulated artificial light at night had a lesser effect. Under dim light, larvae and adults increased expression of the cone opsin genes, *sws2a, rh2c* and *lws*, within a few days and larvae also decreased densities of cones, inner nuclear layer cells and ganglion cells. These changes likely enhanced vision under the altered light conditions. Thus, our study suggests that plasticity mainly comes into play when conditions are extremely different to the species’ natural light environment, *i*.*e*., a diurnal fish in ‘constant night’. Finally, in a rescue experiment on adults, shifts in opsin expression were reverted within 24 hours. Overall, our study showed rapid, reversible light-induced changes in the retina of *A. triostegus*, demonstrating phenotypic plasticity in the visual system of a reef fish throughout life.

## Introduction

Teleost fishes inhabit diverse environments, ranging from coral reefs to the deep sea, and their visual systems have adapted accordingly (Cortesi et al., 2020; de Busserolles et al., 2020; Lythgoe, 1979; Walls, 1942). Some of their best characterised visual adaptations are in the retina. The retina comprises the photoreceptor layer, the inner nuclear layer (INL) and the ganglion cell (GC) layer (Lamb, 2013). Within the photoreceptor layer, rods facilitate scotopic (dim light) vision and contain the rhodopsin protein, RH1 (rhodopsin), while cones facilitate photopic (bright light) vision, and contain several opsins: SWS1 (short wavelength-sensitive 1, ultraviolet), SWS2 (violet-blue), RH2 (medium wavelength-sensitive 2, blue-green) and LWS (long wavelength-sensitive, yellow-red) (Bowmaker, 2008). Visual signals from the photoreceptors are conveyed to the INL, the primary layer for opponent processing (Baden & Osorio, 2019). Finally, the signals are summated in the GC layer, where a trade-off between luminous sensitivity and visual acuity occurs. Generally, lower GC densities enhance sensitivity by increasing the summation of visual signals, while higher GC densities improve acuity by increasing the resolution at which signals are sampled (Collin, 1997; Warrant, 1999).

Changes within the retina can reflect visual adaptations to specific environments. These adaptations may occur at the cellular level, involving the size, number and distribution of retinal cell types (Collin & Shand, 2003; Yoshimatsu et al., 2020), or molecular level, concerning the opsins or other parts of the phototransduction machinery (Carleton et al., 2020). Some of these visual adaptations have emerged over evolutionary timescales. In marine fishes, this is exemplified by differences in spectral sensitivity between species inhabiting various depths [*e*.*g*., shallow vs. deep; (Douglas et al., 2003; Douglas & Partridge, 1997; Schweikert et al., 2018a; Schweikert et al., 2018b)], or habitat types [*e*.*g*., coastal vs. pelagic; (Lythgoe et al., 1994; Marshall et al., 2015)]. However, vision may also be plastic within the lifetime of an individual, such as during its development (Evans & Fernald, 1993; Pankhurst, 1987; Shand, 1997; Shand et al., 2000; Siebeck & Marshall, 2007) and between seasons (Shimmura et al., 2017; Stieb et al., 2016). For example, a shift to a nocturnal lifestyle during ontogeny has been correlated with scotopic remodelling of the retina in reef fishes (Fogg et al., 2022; Shand, 1997). These ontogenetic and seasonal variations are regular, predictable alterations to the light environment that a species will have encountered over many preceding generations, potentially allowing genetic signals to contribute to adaption, such as has been found in killifish (Fuller et al., 2005) and damselfish (Stieb et al., 2016). However, less is known about the ability of fishes to adapt to more irregular and unpredictable changes in the light environment.

The light environment can change unpredictably due to both natural causes (*e*.*g*., weather patterns), and anthropogenic causes [*e*.*g*., light pollution (Davies et al., 2014)]. Therefore, it might be expected that the visual system harbors some degree of phenotypic plasticity to maintain optimal visual performance under these irregular conditions. Indeed, plasticity in opsin gene expression has been observed in several teleost species placed under unpredictably altered light conditions, such as damselfish and cardinalfish (Luehrmann et al., 2018), African cichlids (Dalton et al., 2015; Härer et al., 2017; Irazábal-González et al., 2021; Wright et al., 2020), guppies (Kranz et al., 2018) and Senegalese sole (Frau et al., 2020). In a handful of species, opsin gene expression plasticity was even observed across several life stages [killifish, (Fuller & Claricoates, 2011; Fuller et al., 2010); African cichlids, (Nandamuri et al., 2017b)]. Plasticity in retinal morphology has also been observed, for example, in African cichlids (Karagic et al., 2018; Wagner & Kröger, 2005). In most cases, the plastic responses observed in fishes seem adaptive, with changes to opsin gene expression and retinal morphology likely maximising visual capabilities in the novel light conditions. However, considerable interspecific variation in the responses suggests that, although environment drives the plastic response, phylogeny may constrain it. Likewise, little is known about intraspecific differences in visual plasticity, *e*.*g*., at different life stages. As such, there are major gaps in our understanding of phenotypic plasticity in the visual system, particularly in reef fishes, whose environment continues to change unpredictably due to anthropogenic causes, such as algal blooms and light pollution [reviewed in (Marshall et al., 2019)].

To fill this knowledge gap, we investigated the capacity of the visual system to adapt to stochastic changes in light conditions in both larval and adult stages of a coral reef fish, *Acanthurus triostegus* (convict surgeonfish). This species is widely distributed (Froese & Pauly, 2019) and, unlike for many marine fishes, earlier life stages can be easily and consistently obtained from the wild (Besson et al., 2017). Importantly, the ecology and visual system of this species has been well-characterised. *A. triostegus* is a diurnal species which consumes zooplankton in the upper layers of the open ocean as larvae but rapidly switches to an algal diet on the reef after it metamorphoses into a juvenile (Abitia et al., 2011; Frédérich et al., 2012). This surgeonfish has a well-developed colour vision system to match its diurnal lifestyle, with six opsins (*rh1, sws2a, sws2b, rh2a, rh2c* and *lws*) expressed in settlement larvae and adults (Cortesi, unpublished) and high cone densities in settlement larvae (Besson, 2017).

Here, we exposed settlement larval and adult *A. triostegus* to changed light conditions, *i*.*e*., constant dim light, constant bright light or simulated artificial light at night. A subset of adults was also used for a rescue experiment involving return to a normal light environment after exposure to constant dim light. Following light treatment, the opsin gene expression repertoire in larvae and adults was confirmed using transcriptomics, and opsin gene expression was measured using quantitative PCR. Retinal cell densities were also assessed histologically in larvae. Using this approach, we aimed to contribute to the following unresolved questions relating to the phenotypic plasticity of vision in fishes: 1) Do reef fishes show phenotypic plasticity in the visual system throughout life? 2) Does plasticity occur at both molecular and cellular levels? 3) How rapidly does a plastic response occur?

## Materials and Methods

### Animal collection and care

Details of all animals used in this study are given in Table S1. All individuals were convict surgeonfish, *Acanthurus triostegus* (Linnaeus, 1758). Settlement-stage larvae, larvae that have just transitioned to the reef, were collected at night using hand nets and a crest net positioned on the reef crest of the lagoon at Temae, Moorea, French Polynesia (17°29’S, 149°45’W) in March 2019 (Besson et al., 2020; Lecchini et al., 2004). Adults were collected with clove oil (15% solution) and hand nets on the Great Barrier Reef around Lizard Island, Australia (14°40′S, 145°27′E) in July 2021.

Immediately following collection, fish were transferred to aquariums at the corresponding research station [Centre for Island Research and Environmental Observatory (CRIOBE) on Moorea, and Lizard Island Research Station (LIRS) on Lizard Island]. Animal collection, care and euthanasia followed procedures approved by the University of Queensland Animal Ethics Committee (QBI 304/16). All collections in Australia were conducted under a Great Barrier Reef Marine Park Permit (G17/38160.1) and Queensland General Fisheries Permit (180731) and those in French Polynesia were conducted in accordance with French regulations.

### Light treatment

On the morning following collection, individuals at each life stage were euthanised on day zero (D0) as baseline controls and all other fish were transferred into aquaria for light treatments. Larvae were exposed to five light treatments (Figure 1): 1) 12L12D outdoor: an outdoor control tank which received 12 h of natural bright light (*i*.*e*., sunlight) and 12 h of natural dim light (*i*.*e*., moonlight) and was not subject to artificial light, 2) 12L12D indoor: an indoor control tank exposed to 12 h of bright light (*i*.*e*., artificial light at sunlight intensity; 50,000 lux) and 12 h of dim light (*i*.*e*., artificial light at moonlight intensity; 2 lux), 3) 12L12AL: 12 h of bright light and 12 h of artificial light at streetlight intensity (50 lux) to simulate night-time light pollution, 4) 24L: 24 h of bright light, or 5) 24D: 24 h of dim light. Larvae were euthanised on day three (D3) or six (D6) of exposure (gene expression: *n* = 5 – 6, N = 62; histology: *n* = 4, N = 44; numbers include baseline controls; one eye per individual for each analysis). Preliminary analyses suggested that larvae only showed consistent significant changes under 24D. Therefore, in addition to baseline controls, adults were exposed to three light treatments, two similar to the larvae with sampling at D6, plus a rescue treatment (Figure 1). Adult light treatments were: 1) 12L12D indoor control for six days, 2) 24D for six days or 3) a rescue treatment in which fish were exposed to 24D for six days and were then returned to 12L12D indoor for 24 h (gene expression: *n* = 5; N = 20; numbers include baseline controls). All animals were euthanised at 7:30-9:00 am to avoid circadian effects on opsin gene expression (Yourick et al., 2019).

**Figure 1.**
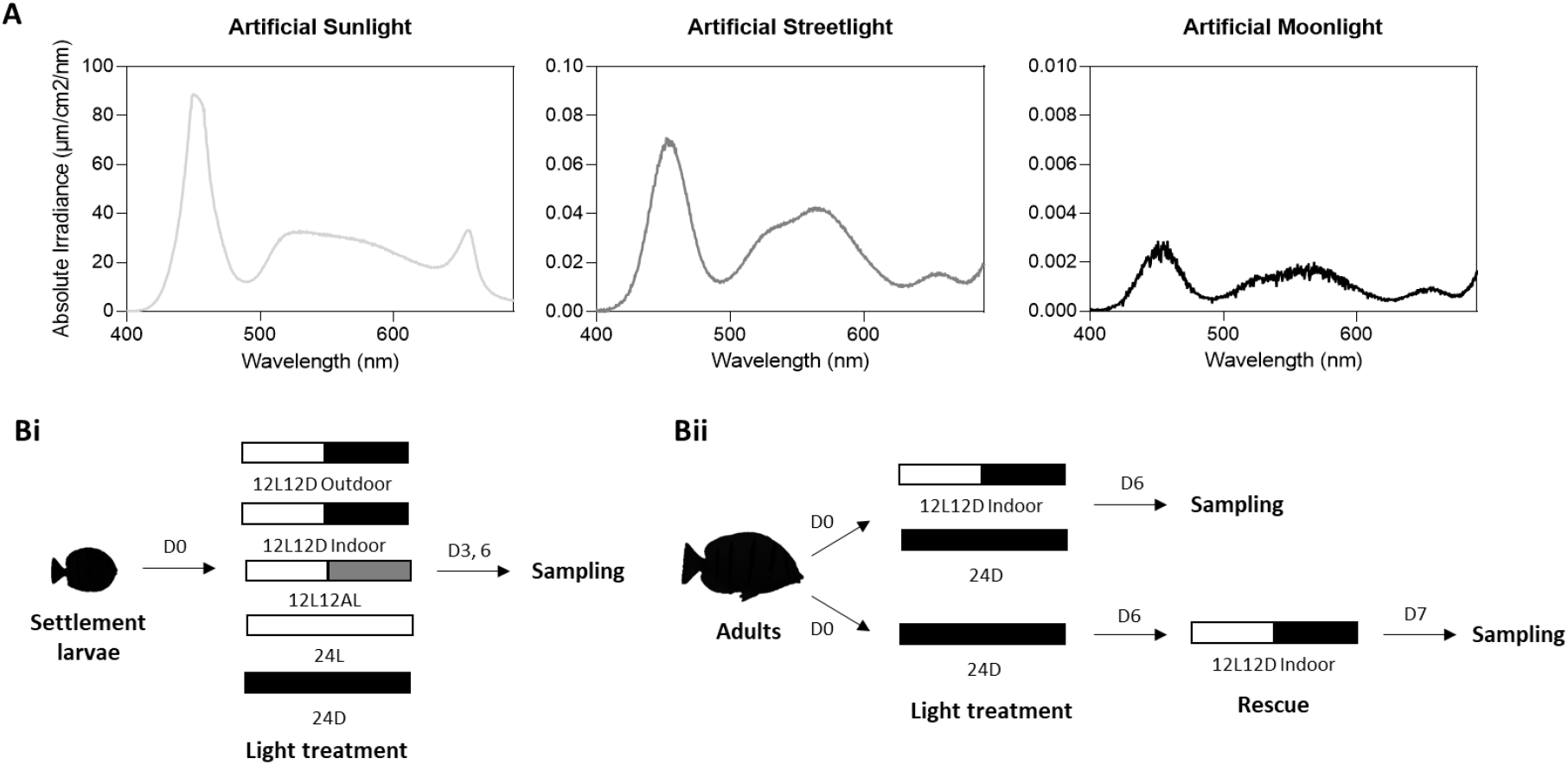
Light treatments. (A) Absolute irradiance (μm/cm^2^/nm) of downwelling light at different wavelengths (nm) for the treatments used in this study. Conditions included artificial light at sunlight, streetlight and moonlight intensity equating to approximately 50,000 lux, 50 lux and 2 lux, respectively. (B) Experimental design for (i) settlement larvae and (ii) adults. Settlement larvae were exposed to five light treatments and then sampled at days (D) 3 and 6. 12L12D outdoor represents exposure to 12 h of natural bright light (*i*.*e*., sunlight) and 12 h of natural dim light (*i*.*e*., moonlight). 12L12D indoor was 12 h each of artificial bright light (*i*.*e*., artificial sunlight) and artificial dim light (*i*.*e*., artificial moonlight). 12L12AL was 12 h of artificial bright light and 12 h of artificial ‘streetlight’. 24L was 24 h of artificial bright light. 24D was 24 h of artificial dim light. Adults were exposed to three light treatments, two similar to the larvae with sampling at D6, plus a rescue treatment. The rescue involved exposure to 24 h of artificial moonlight for six days followed by 24 h in the indoor control treatment (*i*.*e*., 12L12D indoor). Note that D0 baseline controls were also sampled for both stages. L, light (sunlight intensity); D, dark (moonlight intensity); AL, artificial lamplight (streetlight intensity).

Adults were fed twice daily, however, larvae were not fed, as this species likely does not feed during the post-settlement remodelling phase (McCormick et al., 2002; Randall, 1961). Fish were housed in glass tanks coated in black tarp filled with 27-29°C seawater. All artificial light treatments used broad spectrum LEDs (VIPAR spectra, model V165) covered with a diffusion filter (LEE Filters; www.leefilters.com). Different light intensities were achieved by varying lamp brightness settings and adding neutral density filters (0.3 ND, 0.6 ND and 0.9 ND; LEE Filters). The absolute irradiance of downwelling light in the tanks were measured using a 1000 µm optic fibre connected to a Jaz spectrometer and SpectraSuite software (Ocean Optics) (Figure 1).

### Mortality, growth and sample preservation

Fish mortality was recorded upon twice-daily inspection of the aquaria. Following euthanasia at the end of the experiment, all individuals were photographed with a ruler and their standard length (SL) and body height (BH; defined as in Besson (2017)) were measured from the images using Fiji v1.53c (Schindelin et al., 2012). Since altered light conditions can impact morphological growth (Karagic et al., 2018), growth changes were also assessed using body depth (*i*.*e*., BH divided by SL) as it is a good indicator of developmental progress in this species (Besson, 2017; McCormick, 1999).

Following photography, eyes were immediately enucleated, the cornea and lens removed, and the eye cup preserved in either RNAlater (Sigma-Aldrich) or 4% paraformaldehyde [PFA; 4% (w/v) PFA in 0.01M phosphate-buffered saline (PBS), pH 7.4] depending on the analysis (see below for details).

### Transcriptomic assessment of the opsin gene expression repertoire

One transcriptome per time point per condition was sequenced for settlement larvae (*N* = 11) to verify species identity, confirm the opsin gene expression repertoire, and extract housekeeping gene sequences. Firstly, retinal tissue preserved in RNAlater was digested using Proteinase K [New England Biolabs (NEB)], total RNA was isolated using the Monarch Total RNA Miniprep Kit (NEB) and genomic DNA was removed using RNase-free DNase (NEB). Quality and yield of RNA was assessed using the Eukaryotic Total RNA 6000 Nano kit and the Queensland Brain Institute’s Bioanalyser 2100 (Agilent technologies). RNA extractions were shipped on dry ice and retinal transcriptome libraries were prepared from total RNA using the NEBNext Ultra RNA library preparation kit for Illumina (NEB) at Novogene’s sequencing facilities in Hong Kong and Singapore. The concentration of each library was checked using a Qubit dsDNA BR Assay kit (ThermoFisher) prior to barcoding and pooling at equimolar ratios. Libraries were sequenced as 150 bp paired-end reads on a HiSeq 2500 using Illumina’s SBS chemistry version 4.

Libraries were trimmed and *de novo* assembled as described in de Busserolles et al. (2017). Briefly, read quality was assessed using FastQC (v0.72), raw reads were trimmed and filtered using Trimmomatic (v0.36.6) and transcriptomes were *de novo* assembled with Trinity (v2.8.4) using the genomics virtual laboratory on the Galaxy platform [usegalaxy.org; (Afgan et al., 2018)]. Species-specific cytochrome C oxidase subunit I (COI) were downloaded from GenBank (https://www.ncbi.nlm.nih.gov/genbank/). Opsin gene coding sequences (CDS) for *A. triostegus* were provided by Dr Fabio Cortesi (Cortesi, unpublished) and their identity was confirmed by BLASTn (NCBI, Bethesda, MD, https://blast.ncbi.nlm.nih.gov/Blast.cgi) and comparison to the published sequences of a close relative, the spotted unicornfish, *Naso brevirostris* (Tettamanti et al., 2019). All gene extractions and expression analyses were conducted in Geneious Prime v2021.1.1 (Biomatters Ltd).

The species of origin of each retinal transcriptome was confirmed by mapping trinity assembled transcripts back onto the *A. triostegus* COI CDS. Next, the opsin gene expression repertoire was assessed. Opsin gene paralogs were scored on similarity using pairwise/multiple alignments. The similarity score minus one was used as the gene-specific maximum % mismatch threshold for mapping (paired) transcripts back onto the opsin CDS to ensure that reads did not map to multiple paralogs. Proportional expression of each cone opsin gene was estimated by dividing the number of reads mapped to a specific opsin gene by the total number of reads mapped to all cone opsin genes. Cone opsin genes with expression levels estimated to be at least 1% of total cone opsin gene expression, as well as the highly expressed *rh1* gene, were carried forward for quantitative PCR.

Finally, two housekeeping gene CDS, *actb* (*Beta-actin*) and *elf1a* (*Elongation factor 1 alpha*), were manually extracted from the transcriptome for use in normalising opsin gene expression in quantitative PCR (Yourick et al., 2019). Housekeeping gene extractions were performed by mapping filtered paired reads to published CDS of *Oryzias latipes* (*actb*; S74868) or *Amphiprion ocellaris* (*elf1a;* XM_023263215) with medium sensitivity settings. Matching reads were connected by following single nucleotide polymorphisms (SNPs) across genes with continual visual inspection for ambiguity and were extracted as paired mates to mitigate sequence gaps. The consensus of an assembly of these extracted reads was used as the reference for low sensitivity (high accuracy, 100% identity threshold) mapping. Partial CDS extractions were cyclically mapped using the low sensitivity approach to prolong and subsequently remap reads until a complete CDS was obtained. To confirm the identity of each gene, full coding sequences were checked using BLASTn.

### Real-Time Quantitative Reverse Transcription Polymerase Chain Reaction (Real-time qRT-PCR)

Real-time qRT-PCR was conducted as described by Luehrmann et al. (2018) [also see (Stieb et al., 2016) and Yourick et al. (2019)]. Briefly, total retinal RNA was reverse transcribed into cDNA using the High-Capacity RNA-to-cDNA kit (Applied Biosystems) and visualised using SYBR Green (master (Rox) dye; Roche) on a StepOnePlus Real-Time PCR System (Applied Biosystems). Opsin gene expression relative to two housekeeping (2HK) genes was calculated using the formula below, where *f*_gene_,2HK is opsin gene expression normalised to 2HK gene expression, *E*_house_ and *E*_gene_ are the gene-specific primer efficiencies for HK and opsin genes, respectively, and *Ct*_*house*_ and *Ct*_*gene*_ are the critical cycle numbers for the HK and opsin genes, respectively.

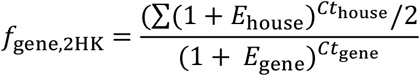

Unique species-specific primers were designed from the opsin and housekeeping gene CDS. Primers produced short (85-100 bp) or long (500-700 bp) amplicons for each gene for quantitative PCR and primer efficiency testing, respectively. To exclusively amplify cDNA, the forward or reverse primer spanned an exon-exon boundary (except for the intronless *rh1*). Primer efficiencies were tested using a three orders of magnitude dilution series of a species-specific opsin pool. To generate the opsin pool, each gene was amplified from cDNA, purified from an agarose gel using the QIAquick PCR purification kit (QIAGEN), quantified with the Agilent 2100 Bioanalyser High Sensitivity DNA kit and mixed in equimolar ratios. All primer details are provided in Table S2. All experiments had three technical replicates with random assignment of samples to each qPCR plate.

### Retinal histology

Retinal histology was conducted on PFA-fixed eyes from settlement larvae. To account for intraretinal variability (de Busserolles et al., 2021), a small piece of tissue was dissected from two different (dorsal and ventral) retinal regions. Tissue was post-fixed in 2.5% glutaraldehyde and 2% osmium tetroxide, dehydrated in ethanol and acetone, and embedded in EPON resin (ProSciTech) using a BioWave Pro tissue processor (PELCO). Radial 1 μm-thick sections were cut on a Leica ultramicrotome (Ultracut UC6) and stained with 0.5% toluidine blue and 0.5% borax. Brightfield images were captured under a 63X objective (oil, 1.4 numerical aperture, 0.19 mm working distance, 0.102 μm/pixel) on a Zeiss Axio upright microscope (Imager Z1).

Since continuous exposure to high-intensity light can cause retinal degeneration (Bernardos et al., 2007; Vera & Migaud, 2009), retinas of fish exposed to 24L were inspected for signs of degeneration prior to analysis. Subsequently, retinal cell densities were estimated from transverse retinal sections as described previously (Fogg et al., 2022; Shand, 1997). Briefly, in Fiji, retinal images were cropped to obtain 100 μm-wide strips, the number of cone outer segments (OS), outer nuclear layer (ONL) nuclei, INL nuclei and GC layer nuclei were counted for three sections per sample using the cell counter plugin and the density of each cell type per 0.01 mm^2^ of retina was calculated. Rod densities were calculated as the difference between ONL nuclei and cone OS densities (Shand, 1994a). Graphs throughout the study were generated using GraphPad Prism software v8.3.1 (www.graphpad.com).

## Results

### Opsin gene expression plasticity in larvae

Opsin gene expression was quantified in larval *A. triostegus* after light treatment (Figure 2). Firstly, larvae expressed the rod opsin, *rh1*, and five cone opsin genes, *sws2a, sws2b, rh2a, rh2c* and *lws*. Under natural light conditions (12L12D outdoor), only the *rh2s* showed a net increase in expression between D0 and D6 (Figure S2; *rh2a*, ns and *rh2c, p*<0.05 at D6), while the other opsins showed minimal net change. Notably, expression differed between the two controls (*i*.*e*., 12L12D outdoor vs. indoor) at D6, with expression higher in the outdoor control for almost all cone opsin genes (*rh2a, p*<0.0001; *sws2a and rh2c, p*<0.05; *lws, p*<0.01). Thus, the artificial light treatments were statistically compared to the age-matched indoor control which used the same light source.

**Figure 2.**
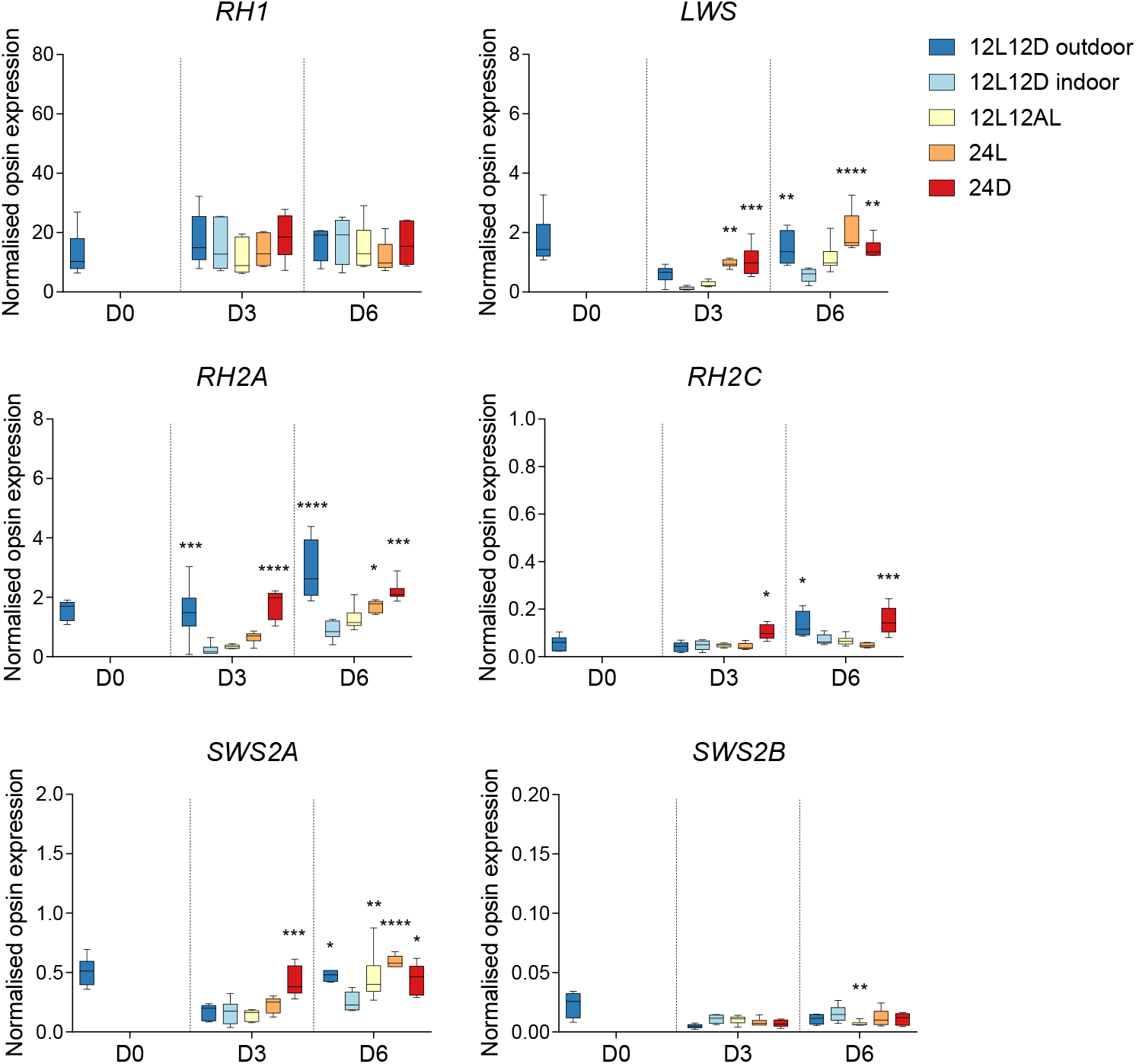
Opsin gene expression in developing fishes under altered light conditions. Opsin gene expression was normalised to two housekeeping genes and time points were taken at days (D) 0, 3 and 6 of exposure to altered light treatments for settlement larvae of *A. triostegus* (*n* = 5 – 6; *N* = 62). Data are mean ± s.e.m. Statistical significance compared to the age-matched 12L12D indoor control (calculated from a two-way ANOVA with Dunnett’s multiple comparisons test): *, p < 0.05; **, p < 0.01; ***, p < 0.001; ****, p < 0.0001. *rh1, rhodopsin-like middle-wavelength sensitive 1* (rod opsin); *rh2, rhodopsin-like middle-wavelength sensitive 2; sws2, short-wavelength-sensitive 2; lws, long-wavelength-sensitive*.

Under all artificial light treatments, changes to opsin gene expression were limited to the cone opsin genes, with no changes to *rh1* expression (Figure 2). The most consistent changes occurred under 24D, in which the dominantly expressed cone opsin genes showed increased expression (*sws2a, p*<0.05; *rh2a* and *rh2c, p*<0.001; *lws, p*<0.01). This trend was evident at both D3 and D6. Notably, these cone opsin genes naturally increased in expression over the experiment (*i*.*e*., in the outdoor control), so increased expression levels under 24D compared to the age-matched indoor control resembled an acceleration to a later ontogenetic stage in the former. Additionally, a second trend was observed in a smaller subset of the cone opsins genes (*i*.*e*., *sws2a, rh2a* and *lws*) in response to the brighter artificial light conditions (*i*.*e*., 12L12AL and 24L). Notably, these genes showed similarly high expression levels under 24D and 24L at D6. However, when only the brighter conditions were considered, a trend towards a graded effect was observed. Effectively, gene expression under 24L was higher than under 12L12AL, which was higher than the indoor control. Thus, increasing exposure to artificial light at night resulted in increasing expression of *sws2a, rh2a* and *lws* at both D3 and D6.

### Opsin gene expression plasticity in adults

Opsin gene expression was also quantified in adult *A. triostegus* after light treatment (Figure 3). For the adult experiment, the treatment that elicited the greatest changes in the larval visual system was used (*i*.*e*., 24D), plus the indoor control and a rescue experiment. Firstly, adults expressed the same opsin gene repertoire as larvae, but expressed three of the cone opsin genes, *sws2a, sws2b* and *lws*, at lower levels when comparing D0 baseline controls (Figure S3). Under artificial light treatments, similar results were observed in adults compared to larvae but on a smaller scale, showing trends rather than significant differences. As such, under 24D, there were no major changes in expression for *rh1* or *sws2b*, while a slight increase in *sws2a, rh2c* and *lws* expression was observed. However, in contrast to the larvae, *rh2a* expression did not change under 24D in adults. Finally, the rescue experiment conducted on adults showed that all genes reverted to expression levels comparable to the control (*i*.*e*., D6 - 12L12D indoor) following the rescue (Figure 3). Thus, the 24 h exposure to a normal light regime negated the effects of the 6-day exposure to dim light.

**Figure 3.**
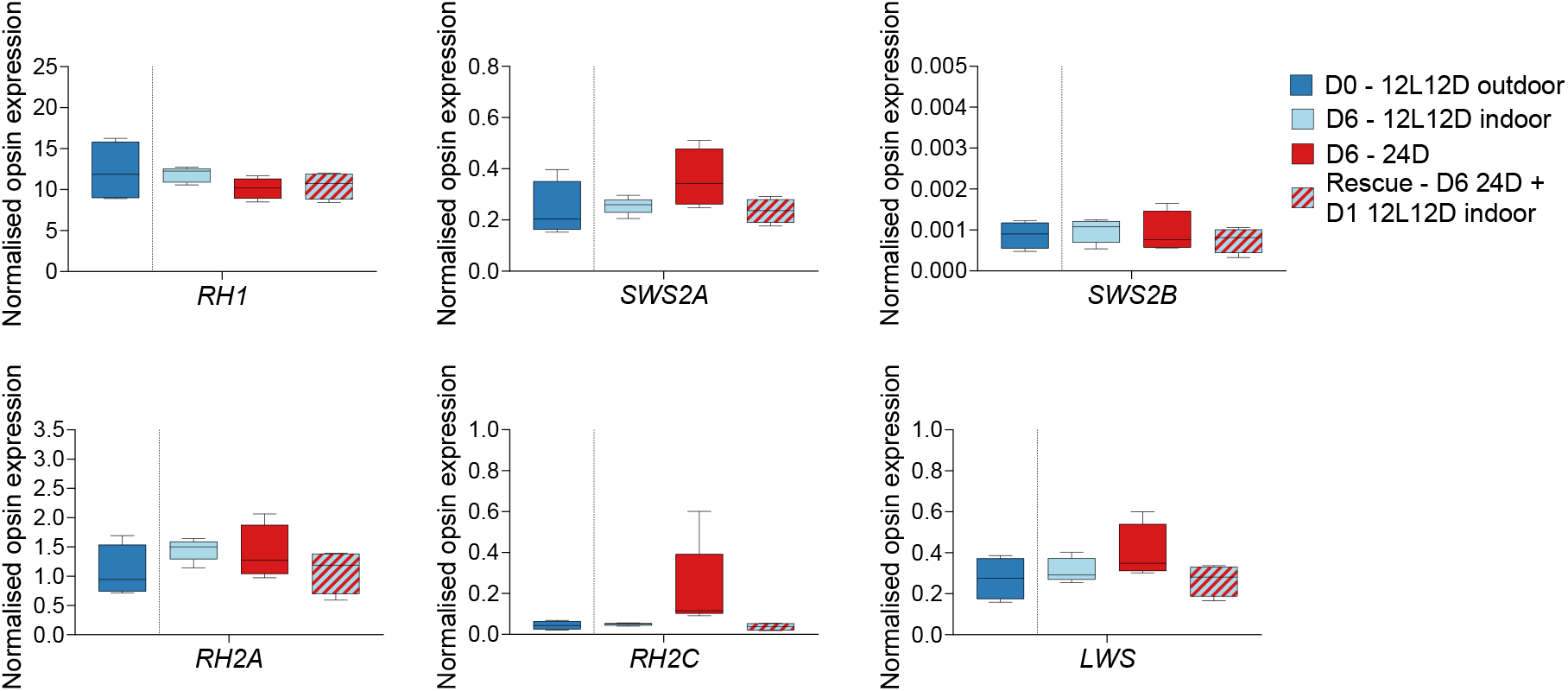
Opsin gene expression in adult fishes under altered light conditions. Opsin gene expression in adult *A. triostegus* was normalised to two housekeeping genes. Time points were taken at days (D) 0 and 6 of exposure to altered light conditions for single-condition exposures and day 7 for the rescue (*n* = 5; *N* = 20). Data are mean ± s.e.m. Statistical significance (calculated from a one-way ANOVA with Kruskal-Wallis multiple comparisons test): no significance detected. *rh1, rhodopsin-like middle-wavelength sensitive 1* (rod opsin); *rh2, rhodopsin-like middle-wavelength sensitive 2; sws2, short-wavelength-sensitive 2; lws, long-wavelength-sensitive*.

### Plasticity in retinal morphology in larvae

Retinal cell densities were also assessed in settlement larvae (Figure 4). Firstly, no overt signs of retinal degeneration were observed under any condition (Figure S1). Secondly, there was no difference between the two control treatments. Furthermore, none of the artificial light conditions altered rod cell densities. Under 12L12AL, a slight decrease in GC densities was observed in the dorsal retina at D6 (*p*<0.05). However, significant density changes across multiple cell types were only observed under 24D. Under 24D, cone densities were lower in the dorsal (D3, *p*<0.0001; D6, *p*<0.01) and ventral retina (D3 and D6, *p*<0.0001) compared to the indoor control at D3 and D6, and INL and GC densities decreased in the dorsal retina at D6 (*p*<0.05).

**Figure 4.**
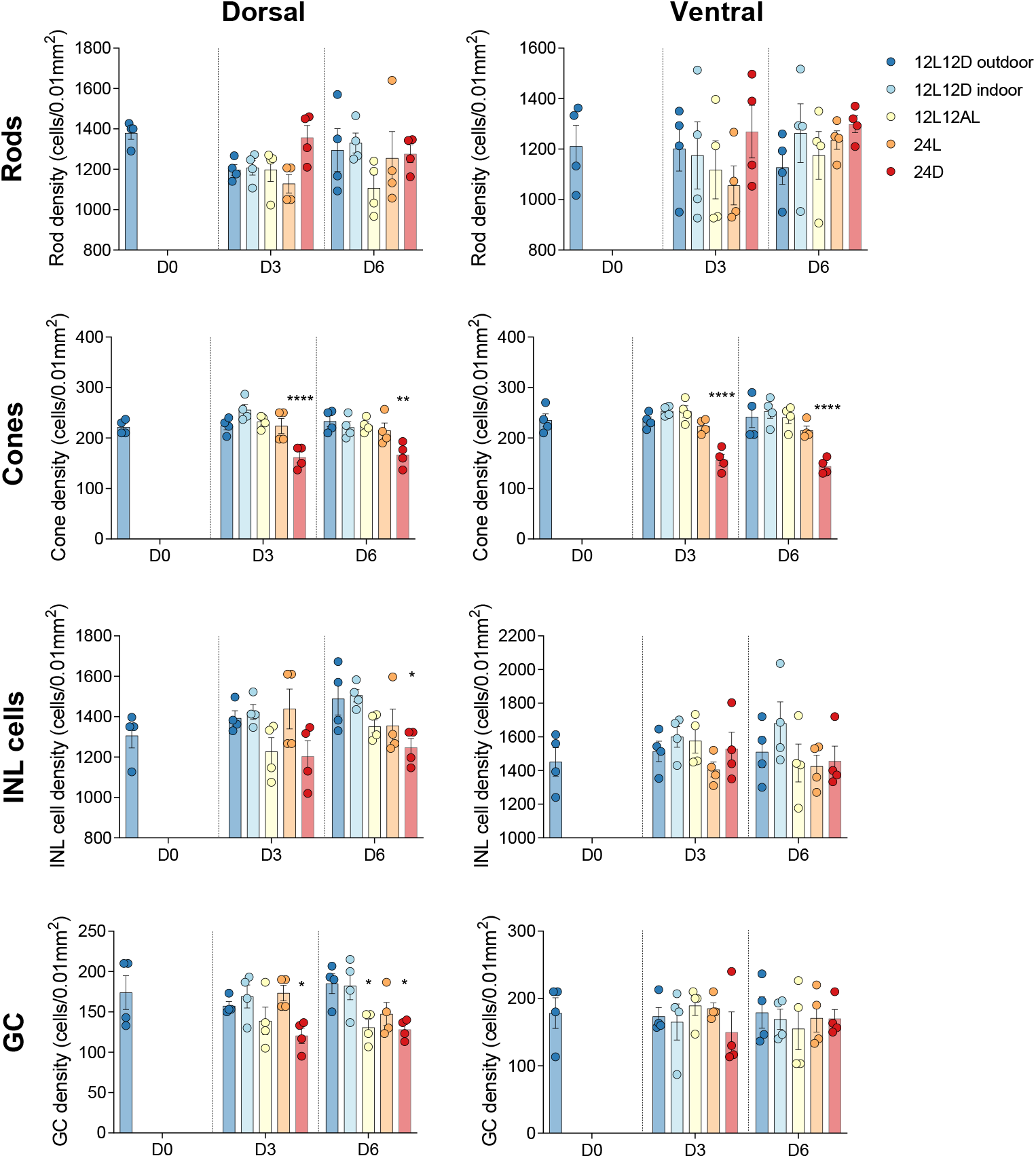
Retinal cell densities in developing fishes under altered light conditions. The densities of rods, cones, inner nuclear layer (INL) cells and ganglion cells (GC) were quantified (as cells per 0.01mm^2^) in the dorsal and ventral retina at days (D) 0, 3 and 6 of exposure to altered light conditions in settlement larvae of *A. triostegus* (*n* = 4; *N* = 44). Data are mean ± s.e.m. Statistical significance when compared to the age-matched 12L12D indoor control (calculated from a two-way ANOVA with Dunnett’s multiple comparisons test): *, p < 0.05; **, p < 0.01; ****, p < 0.0001.

### Body depth and survival in larvae and adults

In both larvae and adults, 100% survival was observed in all treatments. In larvae, body depth decreased under all light conditions, however, the proportional changes were greater between D0 to D3 compared to D3 to D6 (Figure S4; Table S1). Furthermore, body depth reduction differed between the two control conditions, with 14.6% and 18.6% decreases observed in the outdoor and indoor controls, respectively, between D0 and D6. Of the artificial light treatments, 24L induced the most rapid initial decrease in body depth (15.8% by D3 but minimal change thereafter), while 24D induced the greatest overall decrease in body depth (19.1%). Conversely, in adults, no significant changes to body depth were observed (Figure S5; Table S1).

## Discussion

### Ontogenetic tuning of opsin gene expression under natural light

Diurnal reef fish usually express a broad opsin gene repertoire to optimise photopic (bright light) and colour vision [(Cortesi et al., 2016; Luehrmann et al., 2019; Stieb et al., 2019); reviewed in Cortesi et al. (2020)]. The diurnal convict surgeonfish, *A. triostegus*, is no exception, with both larvae and adults expressing six opsins: one rod opsin (*rh1*) and five cone opsins (*sws2a, sws2b, lws, rh2a* and *rh2c*).

The expression of identical opsin gene repertoires at both stages is similar to findings in *N. brevirostris* (Tettamanti et al., 2019), suggesting a direct mode of development in this family. As found in *N. brevirostris*, the opsin genes are differentially expressed between life stages in *A. triostegus*. Ontogenetic changes in opsin gene expression are often related to changes in habitat and diet (Chang et al., 2020; Fogg et al., 2022; Lupše et al., 2021; Shand et al., 2008; Tettamanti et al., 2019). Indeed, *A. triostegus* larvae rapidly increase expression of the green-sensitive cone opsins (*i*.*e*., *rh2a* and *rh2c*) following settlement. This correlates well with a rapid ontogenetic switch to feeding on longer wavelength-reflecting algae after settlement (Abitia et al., 2011; Frédérich et al., 2012) and a switch to a reef environment that is more restricted to mid-range (blue – green) wavelengths compared to the surface waters inhabited by most larvae (Helfman, 2009; Job, 2000). The latter ecological shift is also well aligned with the fact that adult *A. triostegus* expresses lower levels of the cone opsin genes sensitive to wavelengths at the extremities of the light spectrum (*i*.*e*., *sws2a, sws2b* and *lws*).

### Visual plasticity under dim light

Several teleost fishes, including adults of a few reef species, have previously shown visual plasticity under novel light conditions (Härer et al., 2019; Karagic et al., 2018; Luehrmann et al., 2018; Wagner & Kröger, 2005). Here, we demonstrated visual plasticity across ontogeny in a diurnal reef fish, *A. triostegus*. In our study, the strongest responses occurred under constant dim light. This may be because this diurnal species is well-adapted to seeing in photopic conditions (Besson et al., 2020) and therefore, only the dimmer treatments may exceed their natural visual capabilities. Before considering the nature of the visual changes under dim light, it is important to note that the two controls, which used either natural or artificial light, showed differences in opsin gene expression. This finding is congruent with other studies that used aquarium lighting and is likely due to differences in emission spectra (Hofmann et al., 2010; Luehrmann et al., 2018). However, it suggests that the responses in this study may be slightly different to what would be observed on the reef under similar conditions. Regardless, this study permits a controlled investigation of phenotypic plasticity, the nature of which can be reliably interpreted by comparison to the indoor control.

In larvae, constant dim light induced rapid changes at both the molecular and cellular levels. As such, larvae showed increased expression of most cone opsin genes (*i*.*e*., *sws2a, rh2a, rh2c* and *lws*), an acceleration of the developmental progression of cone opsin gene expression, and a decrease in cone, INL cell and GC densities. While increased cone opsin gene expression with decreased cone densities seems contradictory, this finding is consistent with previous work in larval Midas cichlids (Karagic et al., 2018), highlighting our lack of understanding of how opsin gene expression relates to photoreceptor densities. Notably, given the rapid increase in retinal area over settlement (Shand, 1994b), the drop in cone densities found in our study is likely due to a lack of increase in the total number of cones rather than an actual loss of photoreceptor cells. Overall, constant dim light induced rapid changes at both molecular and cellular levels in the larvae.

In adults, constant dim light induced similar changes at the molecular level. As such, adults showed increased expression of most of the cone opsin genes (*i*.*e*., *sws2a, rh2c* and *lws*). However, visual plasticity was less pronounced in adults, potentially due to ongoing development in settlement larvae (Besson, 2017; Fogg et al., 2022; Shand, 1997). Nevertheless, our findings are intriguing since they indicate that *A. triostegus* shows opsin gene expression plasticity in both developing and adult fishes, similar to findings in some freshwater fishes [*e*.*g*., killifish (Fuller & Claricoates, 2011; Fuller et al., 2010) and cichlids (Nandamuri et al., 2017b)]. This emphasises the need for further work on more life stages.

In both life stages, the changes induced by exposure to constant dim light may serve to maximise visual capabilities. At the cellular level, decreased investment in the photopic system (via decreased cone densities) and higher investment in scotopic vision (via increased summation) would theoretically enhance vision in dim light (Pankhurst, 1989; Warrant, 2004). Similarly, at the molecular level, the accelerated developmental progression of cone opsin gene expression may mature visual capabilities to those of later ontogenetic stages which usually inhabit deeper and dimmer waters (Lieske, 1994). Furthermore, increased cone opsin gene expression may increase their capacity to ‘catch’ photons at relevant wavelengths (*i*.*e*., increase their quantum catch) as suggested for other reef fishes (Luehrmann et al., 2018). Although adaptive changes were evident, the changes within the photoreceptors were limited to the cones, as found in damselfishes (Luehrmann et al., 2018). This is intriguing since fishes that naturally inhabit dim-light environments generally adapt their vision via their rods (de Busserolles et al., 2021; de Busserolles & Marshall, 2017). It is possible that as an inherently diurnal species that relies more on photopic vision (Cortesi et al., 2020), *A. triostegus* may have less plasticity in its rods.

### Visual plasticity under bright light

Some teleost fishes have also shown phenotypic plasticity in the retina under increased exposure to bright light [*e*.*g*., Senegalese sole (Frau et al., 2020) and European seabass (Yan et al., 2019)]. Similarly, phenotypic plasticity was evident in larval *A. triostegus* under brighter conditions (24L and 12L12AL). Notably, no retinal degeneration was detected in the current study, likely because it usually occurs at much higher light intensities (Vera & Migaud, 2009). Furthermore, *A. triostegus* showed little to no change in retinal cell densities under the brighter light treatments. This is not so surprising considering this diurnal fish already had well-developed photopic vision (Besson et al., 2020), negating the need for adaptation. Instead, *A. triostegus* showed increased expression of cone opsin genes sensitive to a broad range of wavelengths. This occurred under both constant bright light and simulated artificial light at night (12L12AL), but to a lesser magnitude in the latter. Since photoreceptors undergo bleaching at higher light intensities (Dartnall et al., 1936), this broadly increased expression may represent a compensatory mechanism to help the cones cope with potentially higher levels of opsin turnover.

### Rapid reversion of phenotypic changes

Little is known about whether the visual changes that occur under altered light conditions are dependent on ongoing exposure to the modified conditions. Previous work in zebrafish has shown that a single light-dark transition can rescue clock gene expression in the retina following exposure to constant darkness (Vuilleumier et al., 2006). However, similar studies on opsin gene expression are lacking [but see (Fuller & Claricoates, 2011; Iwanicki et al., 2020)]. Using a rescue experiment on adults, we revealed that shifts in opsin gene expression can be rapidly reverted (within 24 hours) upon return to the control light environment. This represents one of the most rapid cases of light-induced phenotypic plasticity in opsin gene expression reported to date, with previous work showing plasticity in as little as three days in killifish (Fuller & Claricoates, 2011) or a couple of hours in flounders (Iwanicki et al., 2020). Importantly, this finding suggests that visual changes under altered light conditions are reversible in the convict surgeonfish.

### Ecological relevance

Rapid and reversible phenotypic plasticity on a short timescale is likely useful to these fish in a natural ecological context. This is because the light environment that they experience undergoes both diel and seasonal changes. For example, an overcast day can be 100 times dimmer than one with full sunlight (Gaston et al., 2013). Similarly, the duration of daylight is significantly shorter in winter compared to summer (Tseng et al., 2020). Hence, it is not so surprising that *A. triostegus* demonstrates plasticity in response to changes in photoperiod and light intensity. The reversibility of visual changes is also promising in the context of anthropogenic change. The light environment of marine fishes continues to be modified by anthropogenic factors, such as coastal development, commercial fisheries, shipping and tourism (Davies et al., 2013; Davies et al., 2014; Gaston et al., 2017). Therefore, the ability of the visual system to plastically adapt to an anthropogenically modified environment as well as to recover the natural phenotype upon restoration of a natural environment may prove very useful in our changing world.

### Potential mechanisms underlying phenotypic plasticity

Phenotypic plasticity under altered light conditions may be facilitated by various molecular, cellular, or physiological mechanisms. At the physiological level, thyroid hormone signalling is known to play an important role in facilitating developmental remodelling of the retina (Houbrechts et al., 2016), including in *A. triostegus* (Besson et al., 2020). Furthermore, thyroid hormone signalling can mediate shifts in opsin gene expression under constant dim light [*e*.*g*., in cichlids (Karagic et al., 2018)]. However, no differences in thyroid hormone levels have been found between larval *A. triostegus* exposed to altered light conditions compared to natural conditions (O’Connor et al., 2019). Notably, the altered light conditions in this previous study resembled our 12L12AL treatment, which produced minimal visual changes. Thus, thyroid hormone signalling may still be involved in the visual plasticity of *A. triostegus* under more significantly altered light conditions, such as constant dim light.

At the cellular level, increased opsin gene expression in *A. triostegus* could be caused by an increase in outer segment length, as observed in cichlids exposed to constant dim light (Wagner & Kröger, 2005). However, this option seems unlikely in our study given the short timescale. At the molecular level, increased opsin gene expression could result from an increase in packing of the visual pigments in the cone photoreceptors. However, this may also be unlikely given that the density of a visual pigment is thought to be optimised for proper functioning of the light response, so denser packing may disrupt phototransduction (Wen et al., 2009). Instead, co-expression, *i*.*e*., the expression of more than one opsin type in a single photoreceptor (Dalton et al., 2014), may underlie the increase in opsin gene expression, as seen in cichlids under different light environments (Dalton et al., 2015). Co-expression is known to occur in reef fishes (Cortesi et al., 2016; de Busserolles et al., 2021; Stieb et al., 2019), however, further work using in-situ hybridisation will be required to identify co-expression in *A. triostegus*.

Finally, numerous transcription factors and regulatory loci have also been shown to modify opsin gene expression (Nandamuri et al., 2018; Nandamuri et al., 2017a; Sandkam et al., 2020) and mediate morphological changes to the retina (Nelson et al., 2008; Ogawa & Corbo, 2021) in fishes. Therefore, some of these molecular mechanisms may also mediate short-term visual plasticity in *A. triostegus*. Future work looking at differential expression across the retinal transcriptome or, more specifically, at the expression of previously characterised regulatory genes would likely yield some interesting insights.

## Conclusion

The visual system of the convict surgeonfish (*A. triostegus*) showed adaptive phenotypic plasticity across ontogeny, with changes more pronounced in larvae than adults. Specifically, changes under dim light resulted in increased theoretical sensitivity, while those under bright light potentially facilitated increased opsin turnover. However, plasticity was somewhat constrained to light environments which were extremely different to what the fish would naturally experience. Moreover, a rescue experiment on adults showed that shifts in opsin gene expression were rapidly reversible.

These findings enhance our understanding of the capacity of marine fishes to respond to both natural and anthropogenic changes to their environment. This work also brings new questions to light. Firstly, it remains unknown whether the plastic changes would be maintained or would be more pronounced under long-term exposure. For example, would long-term exposure to constant dim light result in a diurnal fish with well-developed scotopic vision? Similarly, although previous studies have suggested interspecific variability in plastic responses (Hofmann et al., 2010), whether the response varies with a species’ ecology (*e*.*g*., diurnal vs. nocturnal) remains unknown. Finally, the mechanisms underlying phenotypic plasticity require further investigation. Thus, future work is required to examine visual plasticity and its underlying mechanisms in a greater breadth of species and over different timeframes.

## Acknowledgements

We acknowledge the Dingaal, Ngurrumungu and Thanhil peoples as traditional owners of the lands and waters of the Lizard Island region from where specimens were collected. We acknowledge the Turrbal/Jagera people as the traditional owners of the land on which the University of Queensland (UQ) is situated, where analyses were conducted. We thank the staff at LIRS and the CRIOBE for support during field work. We thank Robert Sullivan from the Queensland Brain Institute (QBI) Histology Facility and Rumelo Amor from the QBI Advanced Microscopy facility for technical support and advice. We thank the staff at Novogene for library preparation and RNA sequencing. This research was supported by an Australian Research Council DECRA [DE180100949] awarded to F.d.B. F.d.B., F.C. and N.J.M. were supported by the ARC [DE180100949, DE200100620 and FL140100197, respectively]; L.G.F. was supported by the Australian Government and UQ [Research Training Program Stipend] and QBI [Research Higher Degree Top Up Scholarship]; D.L. was supported by the Institute of Coral Reefs of the Pacific (IRCP).

## Author Contributions

L.F., F.d.B., N.J.M. and F.C. designed the study; L.F., D.L., C.G. and F.C. collected the animals; L.F. conducted experiments and collected and analysed the data; F.d.B. and F.C. aided data interpretation; L.F. wrote the initial manuscript. All authors contributed to writing of the manuscript and approved the final version.

## Data Accessibility

Newly identified sequenced and sequenced transcriptomes are available through GenBank and the SRA archive. All other data are available through the UQ eSpace or are provided in the main manuscript or Supplemental Information.

