## Supplemental Information for "Developing and adult reef fish show rapid, reversible light-induced plasticity in their visual system"

### Table of Contents:


| | |
|---|---|
| Table S1. | Page 2 |
| able S2. | Page 4 |
| Figure S1. | Page 5 |
| Figure S2. | Page 6 |
| Figure S3. | Page 7 |
| Figure S4. | Page 8 |
| Figure S5. | Page 9 |

**Table S1. Details of animals used in study. A**ll animals used in this study were convict surgeonfish, *Acanthurus triostegus*. Locations: LI, Lizard Island; MI, Moorea Island. Standard length (SL) and body height are given in millimetres. Time points are given as days of light treatment exposure. Eye: L, left; R, right. Analyses: n.a., only used for morphological measurements; RNA-seq, retinal transcriptome sequenced; qPCR, quantitative PCR used to evaluate opsin gene differential expression; Histology, retinal cell densities quantified.

| Life stage | Location | SL (mm) | Body height (mm) | Body depth | Time point - Light treatment | Eye (molecular) | Eye (histology) | Analyses performed |
|---|---|---|---|---|---|---|---|---|
| Settlement larva | MI | 23.5 | 14.5 | 0.62 | D0 – 12L12D outdoor | R | L | qPCR, Histology |
| | | 25.2 | 15.7 | 0.62 | | L | R | qPCR, Histology |
| | | 26.1 | 14.9 | 0.57 | | R | - | qPCR |
| | | 22.7 | 13.1 | 0.58 | | L | - | qPCR |
| | | 23.1 | 13.4 | 0.58 | | R | L | RNAseq, Histology |
| | | 23.1 | 13.8 | 0.60 | | L | R | qPCR, Histology |
| | | 22 | 14 | 0.64 | | - | - | n.a. |
| | | 24.7 | 12.8 | 0.52 | D3 – 12L12D indoor | R | L | qPCR, Histology |
| | | 26.1 | 12.8 | 0.49 | | L | R | qPCR, Histology |
| | | 27.2 | 13.8 | 0.51 | | R | L | qPCR, Histology |
| | | 25.2 | 12.7 | 0.50 | | - | R | Histology |
| | | 23.9 | 13.3 | 0.56 | | R | - | RNAseq |
| | | 22.9 | 12.5 | 0.55 | | R | - | qPCR |
| | | 23.6 | 12.9 | 0.55 | | - | - | n.a. |
| | | 22.7 | 12.9 | 0.57 | D3 – 12L12AL | L | R | qPCR, Histology |
| | | 26.8 | 13.7 | 0.51 | | R | L | qPCR, Histology |
| | | 26.7 | 13.6 | 0.51 | | L | R | qPCR, RNAseq, Histology |
| | | 25.6 | 12.6 | 0.49 | | R | L | qPCR, Histology |
| | | 26.9 | 14.9 | 0.55 | | - | - | n.a. |
| | | 23.8 | 12.9 | 0.54 | | R | - | qPCR |
| | | 23.9 | 12.1 | 0.51 | | - | - | n.a. |
| | | 24.9 | 12.8 | 0.51 | D3 – 24L | R | L | qPCR, Histology |
| | | 25.7 | 13.3 | 0.52 | | L | R | qPCR, RNAseq, Histology |
| | | 25.1 | 12.6 | 0.50 | | R | L | qPCR, Histology |
| | | 25.3 | 13.1 | 0.52 | | - | R | Histology |
| | | 25.1 | 12.1 | 0.48 | | R | - | qPCR |
| | | 25.8 | 12.3 | 0.48 | | L | - | qPCR |
| | | 25.2 | 13.4 | 0.53 | | R | - | qPCR |
| | | 24.9 | 12.8 | 0.51 | D3 – 24D | L | R | qPCR, Histology |
| | | 25 | 13.2 | 0.53 | | R | L | qPCR, RNAseq, Histology |
| | | 23.7 | 13.1 | 0.55 | | L | R | qPCR, Histology |
| | | 27.4 | 14.2 | 0.52 | | R | L | Histology |
| | | 28.2 | 14.3 | 0.51 | | L | - | qPCR |
| | | 26.1 | 14 | 0.54 | | R | - | qPCR |
| | | 27 | 13.3 | 0.49 | | L | - | qPCR |
| | | 23.4 | 12.9 | 0.55 | D3 – 12L12D outdoor | R | L | qPCR, RNAseq, Histology |
| | | 23.4 | 13.4 | 0.57 | | L | R | qPCR, Histology |
| | | 27.2 | 15.1 | 0.56 | | R | L | qPCR, Histology |
| | | 24.3 | 13.2 | 0.54 | | - | R | Histology |
| | | 27.1 | 14.8 | 0.55 | | R | - | qPCR |
| | | 25.9 | 13.5 | 0.52 | | L | - | qPCR |
| | | 25.9 | 14.1 | 0.54 | | R | - | qPCR |
| | | 24.8 | 11.7 | 0.47 | | L | R | qPCR, Histology |

| | | | | | | | | |
|---|---|---|---|---|---|---|---|---|
| | | 25.5 | 12.2 | 0.48 | D6 – 12L12D indoor | R | L | qPCR, Histology |
| | | 24.9 | 11.5 | 0.46 | | - | R | Histology |
| | | 23.6 | 12 | 0.51 | | R | L | qPCR, RNAseq, Histology |
| | | 25.8 | 13 | 0.50 | | L | - | qPCR |
| | | 25.6 | 12.8 | 0.50 | | R | - | qPCR |
| | | 25 | 12.6 | 0.50 | | L | - | qPCR |
| | | 25.4 | 12.7 | 0.50 | D6 – 12L12AL | R | L | qPCR, Histology |
| | | 24.9 | 12.8 | 0.51 | | L | R | qPCR, Histology |
| | | 25.3 | 12.3 | 0.49 | | R | L | qPCR, Histology |
| | | 24.2 | 12.6 | 0.52 | | L | R | qPCR, Histology |
| | | 25.2 | 12.5 | 0.50 | | R | - | qPCR, RNAseq |
| | | 25.4 | 13.2 | 0.52 | | L | - | qPCR |
| | | 24.4 | 12.4 | 0.51 | D6 – 24L | L | R | qPCR, Histology |
| | | 24.1 | 12.7 | 0.53 | | R | - | qPCR |
| | | 22.9 | 12 | 0.52 | | L | R | Histology |
| | | 24 | 12.3 | 0.51 | | R | L | RNAseq, Histology |
| | | 23.2 | 11.4 | 0.49 | | L | R | qPCR, Histology |
| | | 23.9 | 11.7 | 0.49 | | R | - | qPCR |
| | | 23.9 | 12.2 | 0.51 | | L | - | qPCR |
| | | 25 | 11.7 | 0.47 | D6 – 24D | R | L | qPCR, Histology |
| | | 27.8 | 13.9 | 0.50 | | L | R | qPCR, Histology |
| | | 24.6 | 11.7 | 0.48 | | R | L | Histology |
| | | 25.5 | 11.9 | 0.47 | | L | R | qPCR, Histology |
| | | 24.8 | 12.5 | 0.50 | | R | - | qPCR |
| | | 25.9 | 12.7 | 0.49 | | L | - | qPCR, RNAseq |
| | | 25 | 12.3 | 0.49 | | R | - | qPCR |
| | | 25 | 12.9 | 0.52 | D6 – 12L12D outdoor | L | R | qPCR, Histology |
| | | 24.1 | 12.6 | 0.52 | | R | L | qPCR, Histology |
| | | 25.3 | 12.5 | 0.49 | | L | R | RNAseq, Histology |
| | | 24.2 | 12.8 | 0.53 | | R | L | qPCR, Histology |
| | | 24.6 | 12.7 | 0.52 | | L | - | qPCR |
| | | 25.2 | 12.6 | 0.50 | | R | - | qPCR |
| Adult | LI | 130.2 | 66.7 | 0.51 | D0 – 12L12D outdoor | L | - | qPCR |
| | | 144.5 | 71.1 | 0.49 | | R | - | qPCR |
| | | 106.1 | 55.0 | 0.52 | | L | - | qPCR |
| | | 106.5 | 54.4 | 0.51 | | R | - | qPCR |
| | | 128.6 | 63.1 | 0.49 | | L | - | qPCR |
| | | 91.7 | 44.1 | 0.48 | D6 – 24D | L | - | qPCR |
| | | 131.4 | 68.6 | 0.52 | | R | - | qPCR |
| | | 139.7 | 67.9 | 0.49 | | L | - | qPCR |
| | | 89.6 | 45.7 | 0.51 | | R | - | qPCR |
| | | 96.1 | 46.5 | 0.48 | | L | - | qPCR |
| | | 98.3 | 49.4 | 0.50 | D6 - 12L12D indoor | L | - | qPCR |
| | | 104.5 | 54.5 | 0.52 | | R | - | qPCR |
| | | 96.7 | 48.0 | 0.50 | | L | - | qPCR |
| | | 100.3 | 51.7 | 0.52 | | R | - | qPCR |
| | | 132.3 | 69.2 | 0.52 | | L | - | qPCR |
| | | 146.8 | 73.7 | 0.50 | Rescue – D6 24D + D1 12L12D indoor | R | - | qPCR |
| | | 140.1 | 69.8 | 0.50 | | L | - | qPCR |
| | | 93.0 | 46.1 | 0.50 | | R | - | qPCR |
| | | 99.1 | 48.9 | 0.49 | | L | - | qPCR |
| | | 125.1 | 60.5 | 0.48 | | R | - | qPCR |

**Table S2. Species-specific opsin and housekeeping gene primer sequences used for real-time qRT-PCR.** Quantification primers generated amplicons 85-100 bp in length and pool primer amplicons were 500-700 bp. The efficiency of each qPCR primer set is given as a percentage. All primers were specifically designed for *Acanthurus triostegus*.

| **Opsin** | **Primer** | **Primer sequence (5' to 3')** | **Efficiency (%)** |
|---|---|---|---|
| **RH1** | RH1_fwd | TGTACACCTCAATGCATGGC | 93 |
| | RH1_rev | AGTGACCAGAGCGCAATTTC | |
| | RH1_pool_fwd | CTGTAGCTAACCTCTTCATGG | n.a. |
| | RH1_pool_rev | GTAGATCATTGGGTTGTAGATGG | |
| **SWS2A** | SWS2A_fwd | GCTCTTTTGCATCCAGATACTTC | 93 |
| | SWS2A_rev | CTGACCATACCACCAAGTGTAG | |
| | SWS2A_pool_fwd | TGGGGAGCACAGTAACCTTC | n.a. |
| | SWS2A_pool_rev | GGCAAAGGGAACAGCGAAG | |
| **SWS2B** | SWS2B_fwd | TGTTGGCGGTATGGTAAGC | 91 |
| | SWS2B_rev | GTGAGGCTTGAAGATAAAGTTCC | |
| | SWS2B_pool_fwd | TGTTCTTTCTATTTGTGGCAGG | n.a. |
| | SWS2B_pool_rev | TTCAGCACGAAAAGCAGC | |
| **RH2A** | RH2A_fwd | TCACCATCACTATCACATCCG | 90 |
| | RH2A_rev | GCAACTTCACCTCCTAATGTG | |
| | RH2A_pool_fwd | TGGCAGATCCCATCATGTTC | n.a. |
| | RH2A_pool_rev | GAAAATGAGGAAGACGGGAATG | |
| **RH2C** | RH2C_fwd | GATTCACCATAACCTTCATTACTGC | 93 |
| | RH2C_rev | CTGACCTCCCAAAGTTGCC | |
| | RH2C_pool_fwd | GGAGGAAATGAGCCCAACG | n.a. |
| | RH2C_pool_rev | AACAGGGAAACAGAAGTGGC | |
| **LWS** | LWS_fwd | CAACAATACCAGAGATCCCTTTG | 92 |
| | LWS_rev | AAACATCCAGATTGTTGCAACG | |
| | LWS_pool_fwd | GGCAGAAGAGTGGGGAAAAC | n.a. |
| | LWS_pool_rev | GATAGCCAGAGGAATGAGACAAC | |
| **ACTB** | ACTB_fwd | CTGTGCTGTCTGGAGGTAC | 91 |
| | ACTB_rev | ATGATCTTGATCTTCATGGTGG | |
| | ACTB_pool_fwd | CGACCTCACAGACTACCTC | n.a. |
| | ACTB_pool_rev | CGGACTCATCGTACTCCTG | |
| **ELF1α** | ELF1α_fwd | GTGACAAGATGGGTTGGTTC | 91 |
| | ELF1α_rev | GCAGGATAGCATCAAGAGC | |
| | ELF1α_pool_fwd | GTGAGTTCGAGGCTGGTATC | n.a. |
| | ELF1α_pool_rev | CTTCAGGCAGAGACTCGTG | |

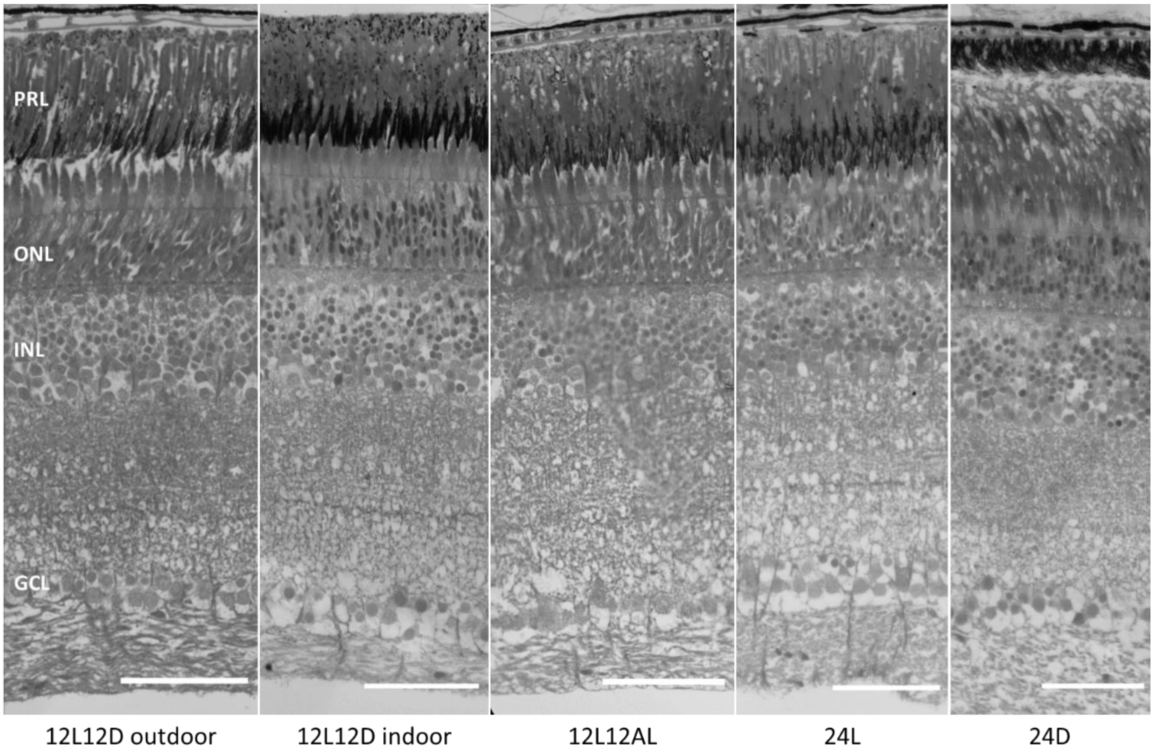


**Figure S1. Retinal morphology under different light conditions.** Representative radial retinal sections from settlement larvae exposed to different light conditions: 12L12D outdoor, 12L12D indoor, 12L12AL, 24L or 24D. The retinas of fishes exposed to 24L are similar in appearance to the controls and no signs of retinal degeneration due to bright-light exposure are apparent. Also, note that the retinas of fishes exposed to 24D are in the dark-adapted state while the rest are in the light-adapted state, which is the reason for the variation in the position of the cones within the photoreceptor layer (PRL). ONL, outer nuclear layer; INL, inner nuclear layer; GCL, ganglion cell layer. *Scale bars*: 50 μm.

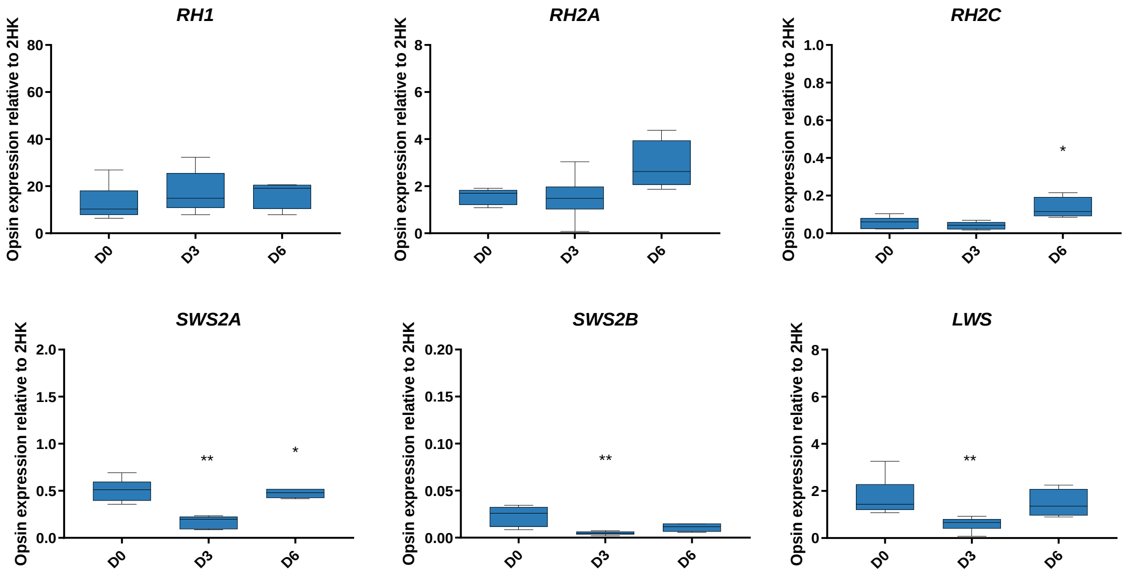


**Figure S2. Opsin gene expression under natural light in developing *Acanthurus triostegus*.** Opsin gene expression was normalised to two housekeeping genes (2HK) and time points were sampled at day (D) 0, 3 and 6 after exposure of settlement larvae to natural light conditions (*i.e.*, 12L12D outdoor) (n = 5 – 6; N = 17). Data are mean ± s.e.m. Statistical significance compared to the preceding time point (calculated from a one-way ANOVA with Kruskal-Wallis multiple comparisons test): *, $p < 0.05$; **, $p < 0.01$.

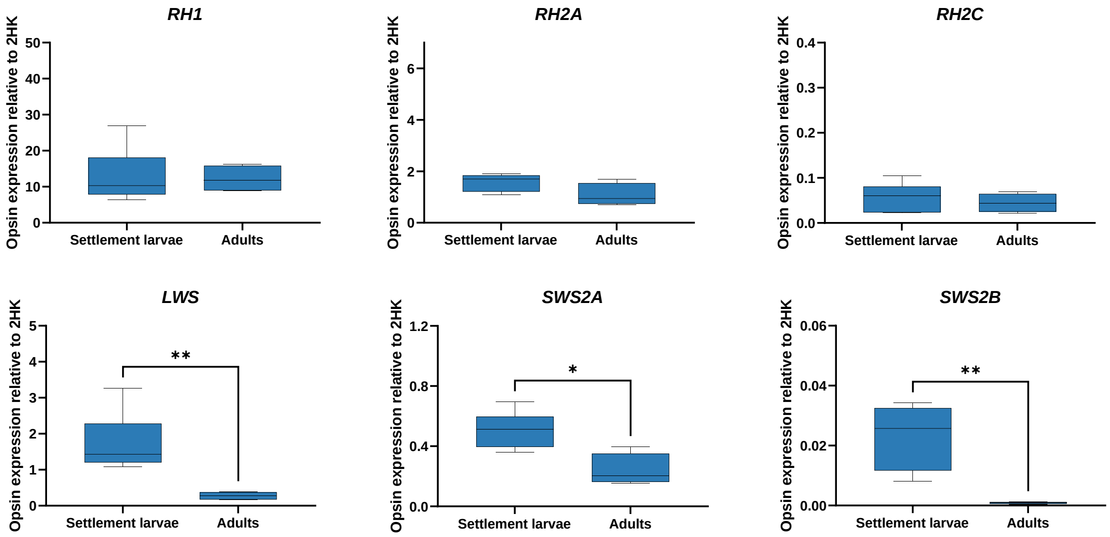


**Figure S3. Changes to opsin gene expression across ontogeny under natural light conditions in *Acanthurus triostegus*.** Opsin gene expression was normalised to two housekeeping genes (2HK) and data are from wild-caught settlement larvae and adults (*i.e.*, D0 12L12D outdoor) (n = 5 – 6; N = 11). Data are mean ± s.e.m. Statistical significance is for comparisons between life stages (calculated from Mann-Whitney U tests): *, $p < 0.05$; **, $p < 0.01$.

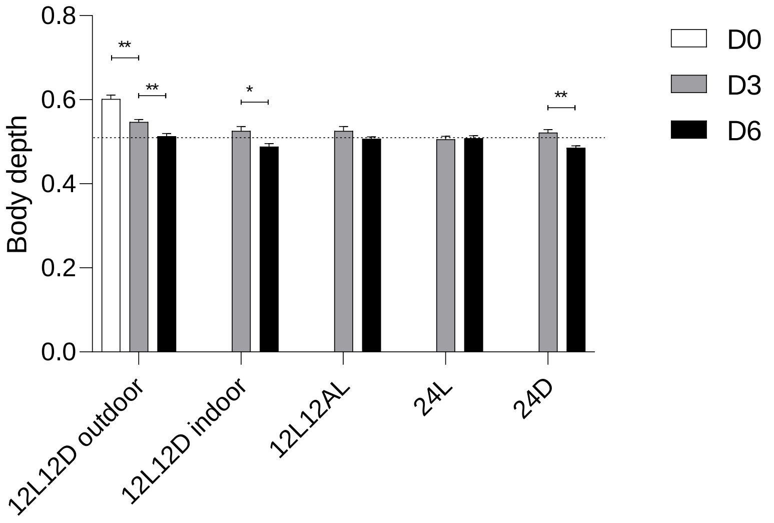


**Figure S4. Body depth under different light conditions in developing *Acanthurus triostegus.*** Body depth was calculated as body height divided by standard length. Time points were taken at day (D) 0, 3 and 6 of exposure to different light conditions for settlement larvae (n = 6 – 7; N = 75). Data are mean ± s.e.m. Statistical significance is a comparison between consecutive time points within the same condition (calculated from Mann Whitney U tests): *, $p < 0.05$; **, $p < 0.01$.

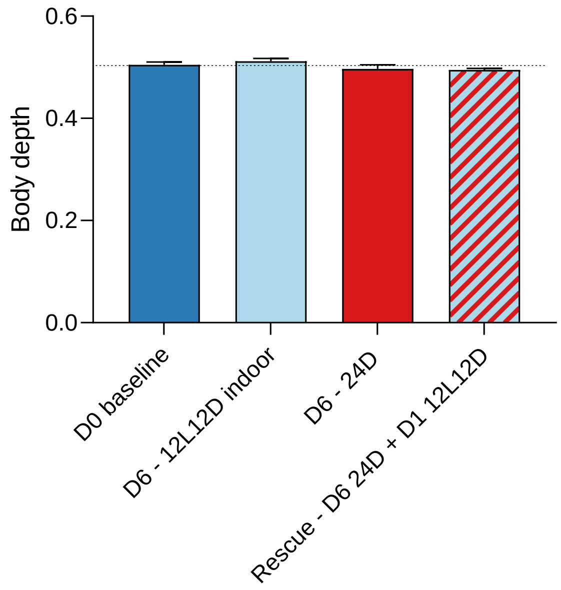


**Figure S5. Body depth under different light conditions in adult *Acanthurus triostegus*.** Body depth was calculated as body height divided by standard length. Time points were taken after adults were exposed to altered light conditions for 0 or 6 days (D), as well as a rescue condition which took 7 days (D) (n = 5; N = 20). Data are mean ± s.e.m. No statistical significance was detected between time points or conditions (calculated from a one-way ANOVA with Kruskal-Wallis multiple comparisons test).
